# Standardizing RNA-seq Analysis of Fungal Pathogens Using BRC-Analytics and Agentic AI: A *Candidozyma auris* Case Study

**DOI:** 10.64898/2025.12.30.697050

**Authors:** Anton Nekrutenko, Danielle Callan, Marius Van Den Beek, Dannon Baker, David Rogers, Aysam Guerler, John Chilton, Hiram Clawson, Scott Cain, Teresa O’Meara, Kelsey Beavers, Michael Schatz, Maximilian Haeussler, Bjorn Gruning, Jeremy Goecks, Sergei Kosakovsky Pond

**Affiliations:** Dept. of Biochemistry and Molecular Biology, The Pennsylvania State University, University Park, PA, USA; Dept. of Biology, Temple University, Philadelphia, PA, USA; Dept. of Biology, Johns Hopkins University, Baltimore, MD, USA; Clever Canary, LLC, Santa Cruz, CA, USA; Baskin School of Engineering, University of California, Santa Cruz, USA; Dept. of Microbiology and Immunology, University of Michigan, Ann Arbor, MI, USA; Texas Advanced Computing Center, The University of Texas, Austin, TX, USA; Dept. of Bioinformatics, Albert-Ludwigs-University Freiburg, Freiburg, Baden-Württemberg, Germany; Moffitt Cancer Center, Tampa, FL, USA

## Abstract

*Candidozyma auris* has emerged as a critical global health threat due to multidrug resistance and healthcare-associated transmission. While RNA-seq has become the primary tool for studying *C. auris* pathogenesis, inconsistent use of reference genomes and bioinformatics tools complicate cross-study comparisons. Here we demonstrate how BRC-Analytics, a platform for pathogen genomics, combined with an agentic AI assistant, enables reproducible RNA-seq analysis. By re-analyzing data from two publications we achieved near-perfect correlation with published results despite annotation version differences. We addressed provenance challenges associated with using AI agents with Galaxy by forcing them to invoke Galaxy’s native tools rather than manipulating data directly. For custom analyses outside Galaxy’s toolset, we provide standalone JupyterLite notebooks that reproduce our analysis without AI involvement. This framework—combining AI-assisted automation with rigorous provenance tracking—establishes a template for standardized, reproducible fungal pathogen genomics. To the best of our knowledge, this is the first example of integration between public data repositories, reproducible analysis workflows, and agentic AI tools. Our subsequent efforts will focus on improving the seamlessness of this integration.

## Introduction

*Candidozyma auris* (formerly *Candida auris*; NCBI:txid498019) represents one of the most urgent antimicrobial resistance threats facing global health systems. First isolated from external ear canal of Japanese hospital patient in 2009 [1], this fungal pathogen has since spread worldwide. CDC classifies *C. auris* as an urgent threat—the first fungal pathogen to receive this designation—due to multidrug resistance (often to all major antifungal classes), healthcare-associated transmission, and 30-60% mortality rates [2,3]. *C. auris* persists on surfaces, colonizes skin, and forms biofilms on medical devices, enabling difficult-to-control nosocomial outbreaks [3]. WHO designates *C. auris* as critical-priority fungal pathogen [4], and NIAID has prioritized development of new therapeutics [5].

Compared to other key human pathogens (such as a SARS-CoV-2 or HIV, for example) the amount of publicly available sequence data for *C. auris* is modest (Table 1). Two categories of projects account for 98% of all data: whole genome sequencing efforts (WGS) and RNA-seq projects. The WGS data are mostly derived from outbreak surveillance efforts conducted by various state public health agencies (Supp. Table 1). The majority of RNA-seq data on the other hand are produced by academic research labs. This reflects the importance of transcriptomic analyses to understanding the fundamental biology of this pathogen. While whole-genome sequencing dominates by run count (26,201 WGS vs 812 RNA-seq runs; 96.3% vs 3.0%), 64 of 237 *C. auris* BioProjects (27%) are RNA-seq studies. This disparity reflects study design: WGS projects sequence many isolates for outbreak surveillance (average 156 runs/project), whereas RNA-seq examines specific biological conditions (average 13 runs/project). Given RNA-seq accounts for over one-quarter of *C. auris* research projects, standardizing analysis is a critical priority.

**Table 1:**
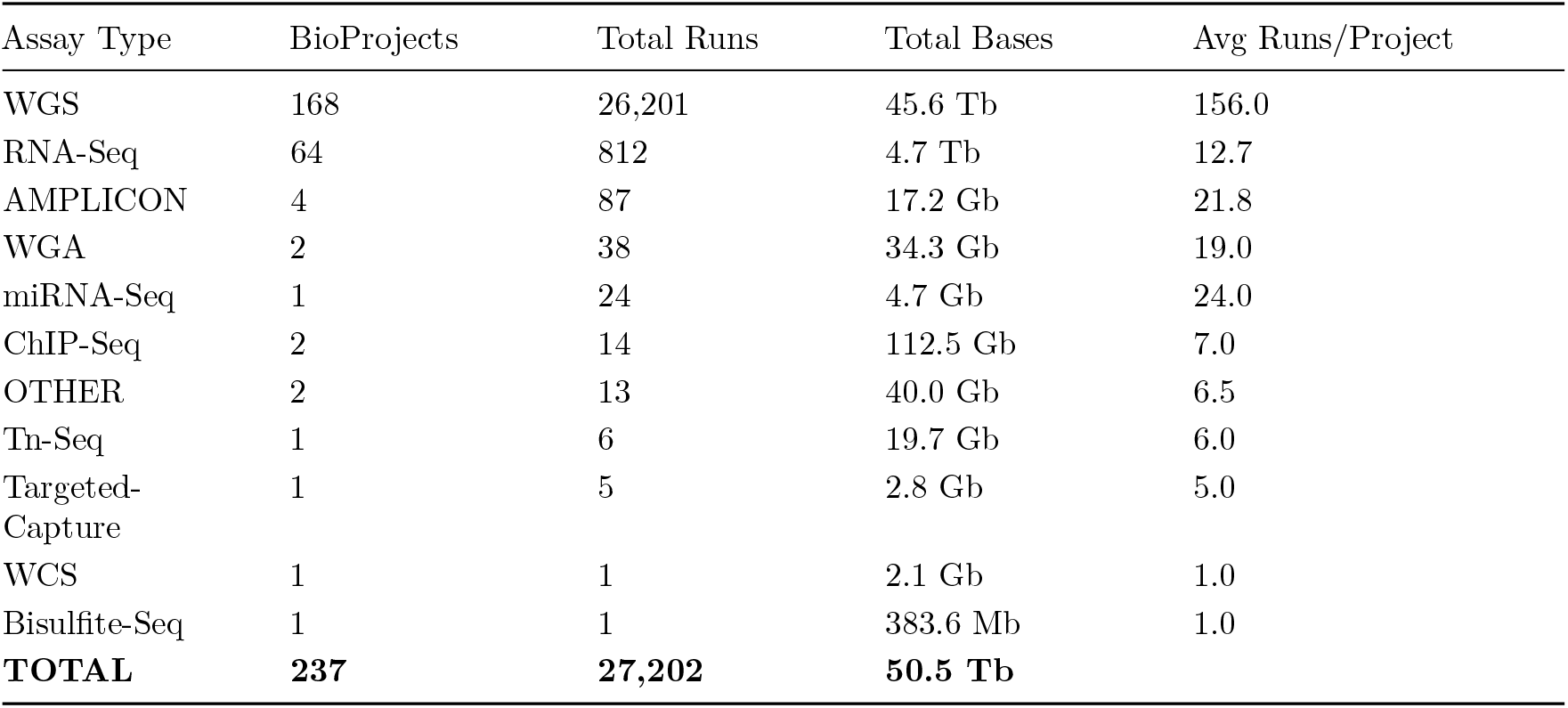
Summary of *C. auris* sequencing data in NCBI SRA (December 2025). BioProject is an NCBI database entry grouping related sequencing runs from a single study. Assay types: WGS = whole genome sequencing; RNA-Seq = transcriptome sequencing; AMPLICON = targeted amplicon sequencing; WGA = whole genome amplification; miRNA-Seq = microRNA sequencing; ChIP-Seq = chromatin immunoprecipitation sequencing; Tn-Seq = transposon insertion sequencing; Targeted-Capture = hybridization capture sequencing; WCS = whole chromosome sequencing; Bisulfite-Seq = DNA methylation sequencing.

To understand the analytical landscape of *C. auris* transcriptomic studies we surveyed all available RNA-seq data associated with that species. Specifically, for all 64 RNA-seq BioProjects listed in Table 1 we attempted to retrieve associated publications. Of 64 BioProjects, 20 (31%) had linked manuscripts (21 papers total, 2018-2025) while 44 remained unpublished or in pre-print stage. For papers with available full text (17/20), we extracted reference genome and analysis tool information (Table 2; also see Supp. Table 2).

**Table 2:** RNA-seq methodology across 20 published *C. auris* studies with linked BioProjects. Numbers in parentheses indicate study count; bracketed numbers are citation references.

| Category | Finding |
| --- | --- |
| **Reference Genome** | B8441/GCA_002759435.x (12/20, 60%); multiple clades (5/20); not specified (2/20) |
| **Alignment Tool** | HISAT2 [6] (7), STAR [7] (5), Bowtie2 [8] (4), BWA [9] (3), TopHat2 [10] (1) |
| **Quantification** | featureCounts [11] (5), HTSeq [12] (4), StringTie [13] (2), Kallisto [14] (2), RSEM [15] (1) |
| **DE Analysis** | DESeq2 [16] (12), edgeR [17] (4), Cufflinks [18] (1) |
| **Publication Years** | 2018 (2), 2021 (4), 2022 (4), 2023 (2), 2024 (5), 2025 (4) |

Despite tool convergence, reference genome usage remains inconsistent. While 60% of published studies use B8441 (GCA_002759435 family), annotation versions vary—some cite only “B8441” without version, others specify GCA_002759435.2 or GCA_002759435.3. This creates reproducibility challenges (e.g., gene identifiers differ between versions) and complicates interpretation of published data in context of new genomes and vice versa. Similarly, tool version reporting is frequently incomplete or absent—papers cite “HISAT2” or “DESeq2” without specifying version numbers, yet algorithm behavior and output can differ substantially between releases. Without precise version information, reproducing published results becomes guesswork. These findings underscore need for standardized platforms specifying precise genome versions, tool versions, and parameters.

Here, we demonstrate how a new environment for the analysis of pathogen, host, and vector data—BRC-Analytics (https://brc-analytics.org)—can be used for standardizing and simplifying RNA-seq analyses using two recent *C. auris* studies as an example. Our approach makes cutting edge tools and powerful computational infrastructure freely accessible to any biologist. Importantly, the combination of BRC-Analytics, the Galaxy platform [19], and Agentic AI tools built on Large Language Models (LLM) tools described here automatically keeps provenance and ensures analytical reproducibility: any analysis conducted within our system can be understood and replicated by others.

## Results

### BRC-Analytics

BRC-Analytics (https://brc-analytics.org) is a browser-based analysis environment designed to make comprehensive and reproducible genomic analyses of infectious diseases accessible to everyone. Developed under the NIAID-funded Bioinformatics Resource Centers (BRCs) program, it leverages the Galaxy platform to enable users to begin with raw sequencing reads and achieve publication-ready results without the need for local software installations or manual data transfers between tools. The platform integrates authoritative genomic data from multiple sources: NCBI Datasets provides reference genomes (currently 5,060 assemblies for 1,920 pathogen, host, and vector taxa, with continuous expansion planned), UCSC Genome Browser supplies genome annotations including gene coordinates and regulatory elements, and EBI ENA facilitates access to public sequence read archive data through local caching for quick searches. BRC-Analytics pairs these data sources with community-curated best-practice analysis workflows covering essential steps like quality control, read mapping, variant identification, and annotation. Galaxy serves not only for launching and running workflows but also as an environment for interpretive analyses through interactive tools like Jupyter. The platform utilizes free cloud-based computation, versioned workflows, and interactive visualizations to create a seamless, reproducible interface. The substantial computational and storage resources required are provided by ACCESS-CI infrastructure in the US, with BRC-Analytics and Galaxy hosted on servers at the Texas Advanced Computing Center (TACC). This approach unifies data and analytical capabilities, making advanced pathogen genomics available to a wider research community.

### Two representative studies

The *Introduction* section above described a survey of all publicly available *C. auris* sequence data with a particular focus on RNA-seq studies and associated publications (Supp. Table 2). From these publications we selected two studies. The first, Santana et al. (2023), identified *SCF1* gene as *C. auris*-specific adhesin essential for biofilm formation and virulence (PRJNA904261) [20]. The second, Wang et al. (2024), showed that glycan-lectin interactions modulate colonization and fungemia (PRJNA1086003) [21]. These two studies are good representatives of *C. auris* RNA-seq methodology. Both use B8441 (Clade I) reference genome, which dominates the field (14/20 published studies). Wang employs `HISAT2/STAR + DESeq2`, the most common pipeline (`DESeq2` in 13/20, HISAT2 in 6/20 studies). Sample sizes of 13 and 6 runs bracket the typical range (median ∼13-15). As 2023-2024 publications, they reflect current practices unlike older studies using outdated tools (`TopHat2, Cufflinks`). Both study adhesion/biofilm phenotypes, the dominant research theme alongside drug resistance.

### The use of Agentic AI

Two aspects of our analysis resist automation: understanding the relationship between datasets deposited to NCBI and actual results described in the two papers and comparison of results produced by us against results in the two manuscripts. Let us look at these challenges in more detail.

The RNA-seq analysis described in this paper is deliberately generic: standard differential expression between two conditions. It can be divided into two parts. The first part produces counts of reads falling within genomic coordinates of each gene. This part is straightforward and is conducted in exactly the same way for all samples. The second part requires reorganizing samples into higher level hierarchy in which they are grouped by conditions that, in turn, contain replicates and so on. When replicating published results, the second part often becomes quite challenging as it requires mapping results described in a paper to actual read-level datasets from NCBI: exactly how samples shown in figure X correspond to dataset Y?

The second challenge is that published data are often analyzed in a context of older (or different) genomic reference. This implies that neither genomic coordinates nor gene ids are matching. Untangling this may be quite complex and may require different approaches depending what specific organism the analysis is performed on.

Both of these challenges may be solved by agentic AI tools powered by LLMs. However, using LLMs through familiar web interfaces—ChatGPT, Claude, or Gemini in a browser—is unsuitable for research problems because artifacts generated during these “chats” are difficult to track, version, or reproduce. Instead, tools like Claude Code (from Anthropic) or Gemini Code Assist (from Google) that operate on a researcher’s own machine are ideal: they keep all artifacts local where they can be preserved and versioned. Crucially, these agents can interact with powerful platforms like Galaxy to execute analyses on robust infrastructure while preserving artifacts necessary for maintaining provenance.

For this analysis we used Claude Code Agent (CCA) produced by Anthropic, configured to interact with the Galaxy platform via an API key that allows CCA to take actions on behalf of the user (see Methods). CCA ran on the local computer used to prepare this manuscript and communicated with Galaxy via Internet (Figure 1A). We are working on integrating agentic AI tools directly into our web platform to avoid managing credentials, eliminating the need to pay for these services, and to improve provenance and reproducibility (Figure 1B; also see Discussion). Before describing our computational setup, we emphasize that any results produced by LLMs must always be verified. For each analysis we first ask CCA to produce a plan of action, review and modify it as necessary, then allow the agent to proceed.

**Figure 1.**
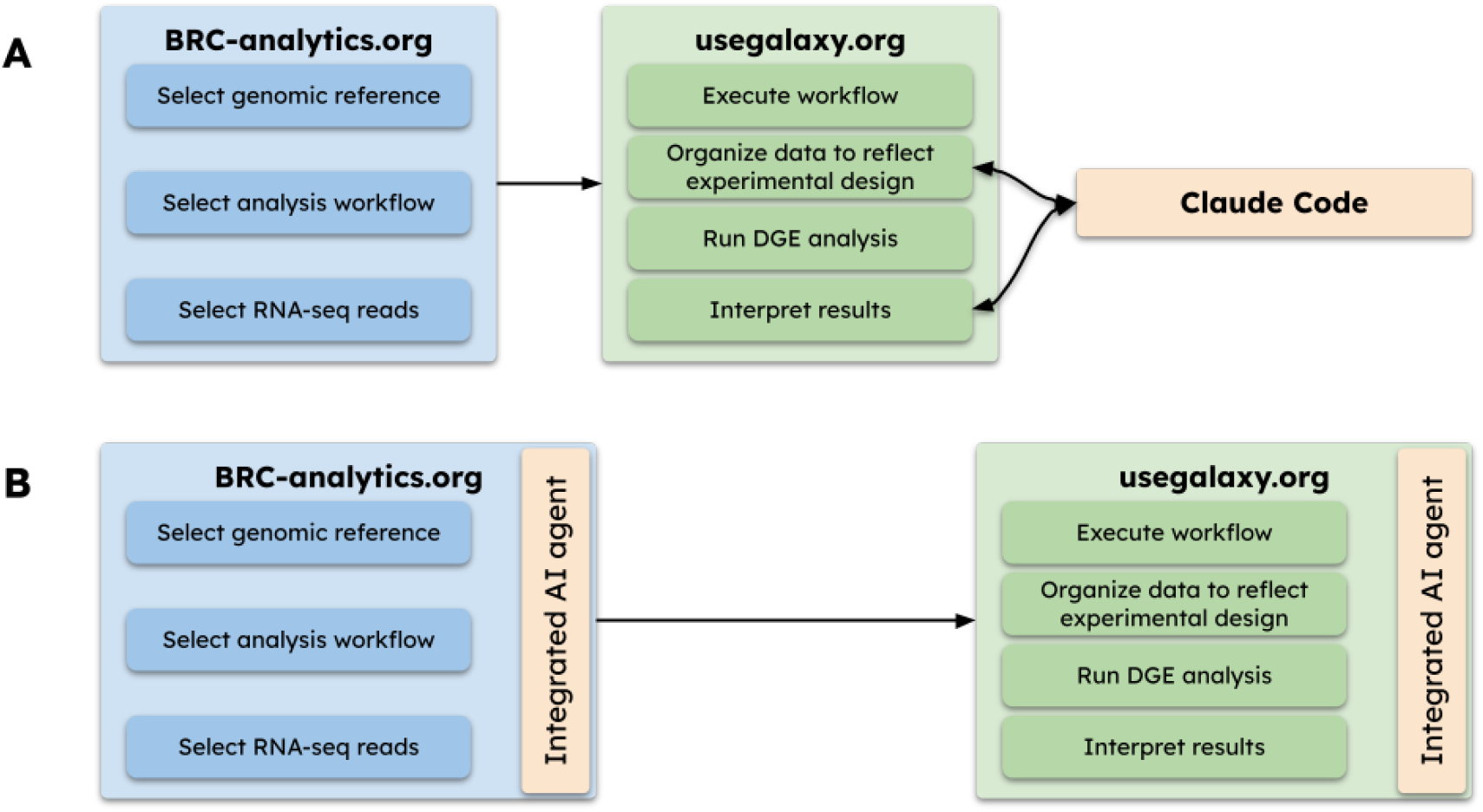
Analysis flow in this manuscript (A) and with integrated AI agents that will be available in future releases (B).

### Organizing data

Authors of the two papers we re-analyze here have deposited sequencing data into NCBI SRA and were given a BioProject identifier—an entity grouping related sequencing runs from a single study. Before performing the differential expression analyses we need to understand how samples deposited to SRA correspond to experimental conditions described in each manuscript. To begin we deposited PDFs of each manuscript along with all available supplemental data into a separate folder on a local computer. We then provided the following instructions to the CCA (we also provide CCA with API key for Galaxy that allows it to access the data and perform operations; for setup see AI agent integration setup within Materials and Methods):

> *I need to split Galaxy dataset collection #244 into several collections corresponding to experimental conditions described in the manuscript (check manuscript pdf and supplemental materials xlsx files in this directory). In order to do this you need to download metadata for sequencing runs for bioproject PRJNA904261 to obtain accessions and metadata. You should then figure out how SRA accessions correspond to experimental conditions described in the paper. You should then present these finding to me, so that I can tell you what to do next*.

This query asks CCA to look at Galaxy dateset collection #244 in history https://usegalaxy.org/u/cartman/h/prjna904261-perm and create a plan for splitting it into three collections corresponding to three strains (conditions) used by Santana et al.: AR0382_WT, AR0387_WT, and tnSWI1. It is important to note that we are not asking for an action. We are asking for a plan that we can review and then decide whether it can be enacted or needs to be modified (see Supplement 1).

In the above prompt we specifically mentioned “Collection #244”—a Galaxy artifact containing read counts for all samples described in this study (can be viewed at https://usegalaxy.org/u/cartman/h/prjna904261-perm). The CCA correctly identified the relationship between datasets and experimental conditions described in the manuscript (Table 3; Supplement 1). After reviewing the plan we instructed CCA to enact it:

**Table 3:** Breakdown of datasets for DESeq2 analysis. For Santana et al. AR0382_WT/tnSWI1 and AR0382_WT/AR0387_WT comparisons were performed. For Wang et al. AR0382 in vitro/AR0387 in vitro and AR0382 in vivo/AR0387 in vivo comparisons were performed.

| Study | Condition | SRR Accessions | Description |
| --- | --- | --- | --- |
| Santana et al. | AR0382_WT | SRR22376031, SRR22376032 | Wild-type reference |
|  | AR0387_WT | SRR22376029, SRR22376030 | Poorly adhesive strain |
|  | tnSWI1 | SRR22376027, SRR22376028 | SWI1 mutant |
| Wang et al. | AR0382 in vitro | SRR28790270, SRR28790272, SRR28790274 | In vitro culture |
|  | AR0387 in vitro | SRR28790276, SRR28790278, SRR28790280 | In vitro culture |
|  | AR0382 in vivo | SRR28791430, SRR28791431, SRR28791432 | In vivo infection |
|  | AR0387 in vivo | SRR28791433, SRR28791434, SRR28791437, SRR28791438 | In vivo infection |

> *Go ahead and execute the plan. Once you are done please add name tags to dataset collection containing data we need to used for DeSeq2 analysis. E*.*g*., *label collections with names tags such as AR0382_WT, AR0387_WT, and tnSWI1*.

This step generated three dataset collections in Galaxy history corresponding to the three conditions described in the paper: AR0382_WT, AR0387_WT, and tnSWI1 (Fig. 2). We then repeated this procedure in a separate Galaxy history containing read count derived from Wang et al. 2024 [21].

**Figure 2.**
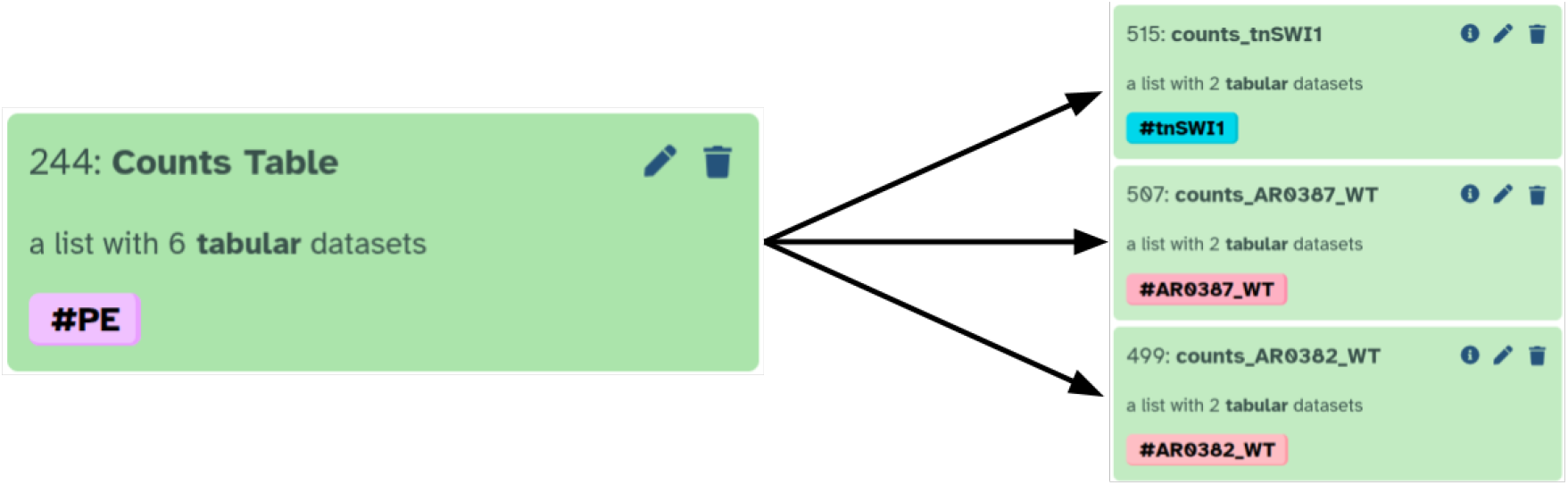
CCA splits collection containing counts into three collections corresponding to three different strains (conditions).

**Figure 3.**
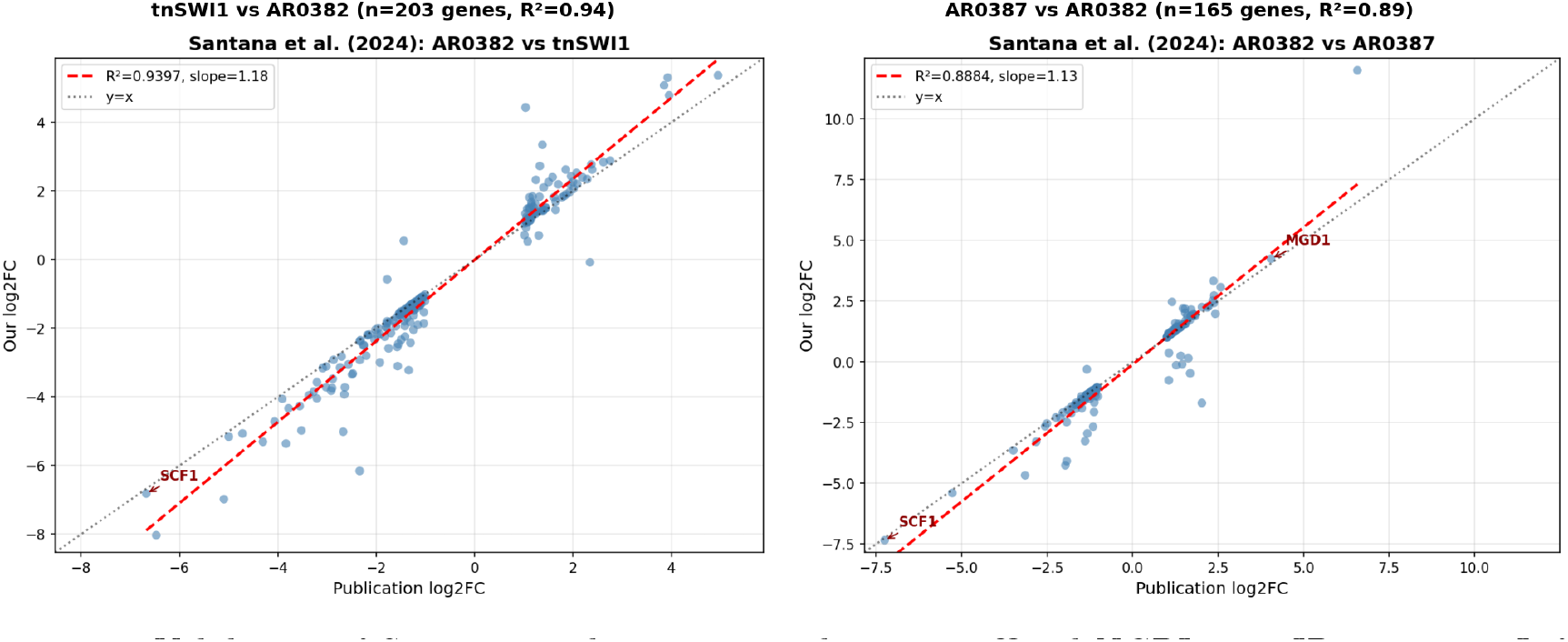
Validation of Santana et al. `DESeq2` results using official NCBI gene ID mapping. Left: tnSWI1 vs AR0382 comparison (n=203 genes, R2=0.94). Right: AR0387 vs AR0382 comparison (n=165 genes, R2=0.89). Red dashed line indicates perfect correlation (y=x). Key gene SCF1 is labeled.

**Figure 4.**
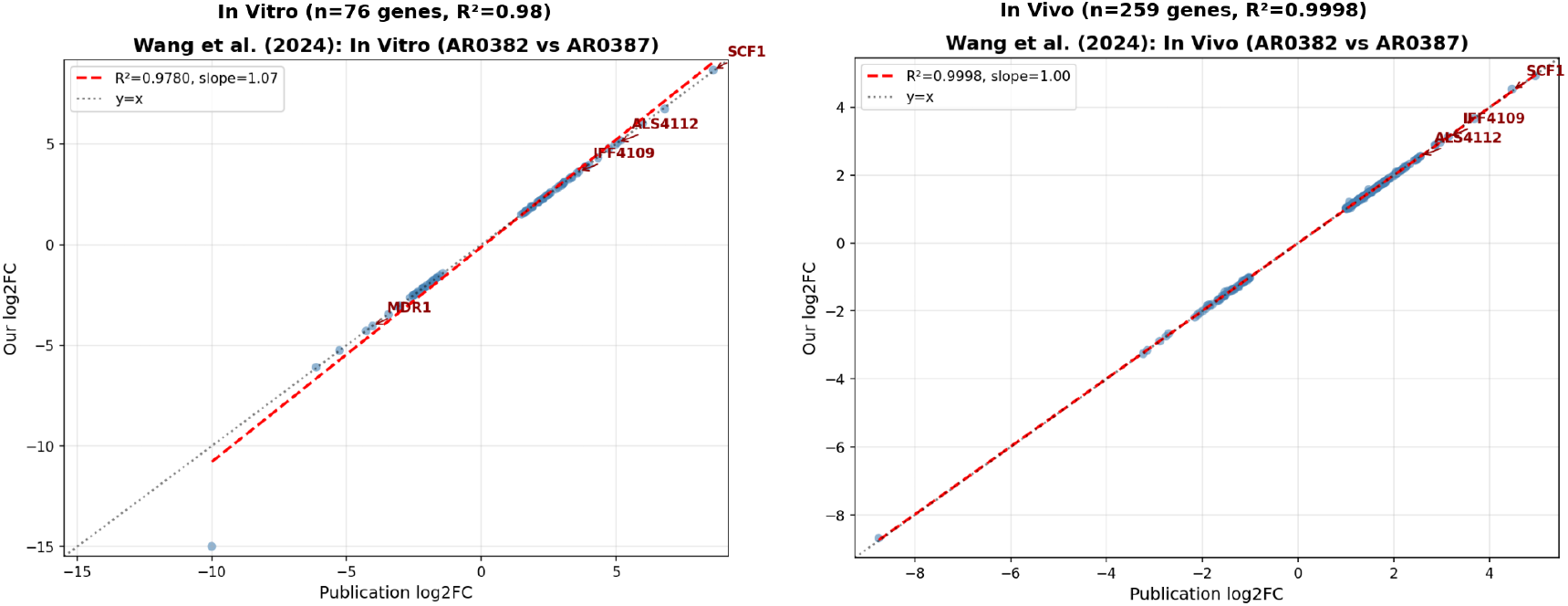
Validation of Wang et al.`DESeq2` results using official NCBI gene ID mapping. Left: In vitro biofilm comparison (n=76 genes, R2=0.98). Right: In vivo mouse catheter model (n=259 genes, R2=0.9998). Red dashed line indicates perfect correlation (y=x). Key adhesin genes SCF1 and ALS4112 are labeled.

### Expression analysis and interpretation

In the previous section we have configured our data so we can perform differential expression. We then re-ran differential expression with `DESeq2` on data from both manuscripts as described in Table 3 and performed a systematic comparison of log2 fold changes against published supplementary data using the following prompt (here, using Santana et al. as an example):

> *Datasets #521 and #523 in https://usegalaxy.org/u/cartman/h/prjna904261-perm represent De-Seq2 results for AR0382_WT/tnSWI1 and AR0382_WT/AR0387_WT comparisons, respectively. Compare them with the results reported in Santana et al. using paper PDF and supplementary files. Use NCBI old_locus_tag attribute for gene ID mapping between annotation versions*.

A technical challenge arose from differences in genome annotation versions. Both published studies used an older *C. auris* annotation with 6-digit gene ID suffixes (e.g., B9J08_001458), while we relied on the latest assembly (GCA_002759435.3) that uses 5-digit suffixes (e.g., B9J08_03708). To reconcile gene identities between versions, we used the official NCBI old_locus_tag attribute present in the GCA_002759435.3 GTF annotation file (this was, in fact, suggested by CCA). This attribute provides authoritative correspondence between old (v2) and new (v3) gene identifiers. We validated this mapping by comparing protein sequences encoded by mapped gene pairs—all pairs showed 100% sequence identity, confirming correct correspondence.

### Comparison with Santana et al. (2023) results

The study compared three strains (Table 3): AR0382_WT, a wild-type highly adhesive Clade I isolate; AR0387_WT, a poorly adhesive clinical isolate; and tnSWI1, a transposon-insertion mutant of AR0382 with disrupted SWI1, a chromatin remodeling factor. Each strain was sequenced in duplicate. The first comparison (Santana et al. Fig. 1D) examined the tnSWI1 mutant versus wild-type AR0382 to identify genes affected by SWI1 disruption. The second comparison (Santana et al. Fig. S5A) contrasted AR0387 against AR0382 to characterize expression differences between adhesive and non-adhesive strains. Both comparisons yielded strong validation metrics. For the first (tnSWI1/AR0382_WT) comparison, we successfully mapped 203 differentially expressed genes and obtained R2 = 0.94 with 99% direction agreement. The second comparison (AR0382_WT/AR0387_WT) mapped 165 genes with R2 = 0.89 and 97% direction agreement. SCF1, the central finding of the Santana study, was the most strongly downregulated gene in both comparisons. The published analysis reported SCF1 (B9J08_001458) with log2 fold changes of -6.68 (Santana et al. Fig. 1D) and -7.25 (Santana et al. Fig. S5A). Our reanalysis identified the corresponding gene (B9J08_03708) with log2 fold changes of -6.82 and -7.35, respectively, confirming the paper’s key finding with minimal deviation.

### Comparison with Wang et al. (2024) results

This study compared two strains with distinct aggregation phenotypes: AR0382 (B11109), a highly aggregative biofilm-forming strain, and AR0387 (B8441), a non-aggregative strain. RNA-seq was performed under two conditions: in vitro biofilm growth (3 replicates per strain) and in vivo colonization of mouse jugular vein catheters (3 replicates for AR0382, 4 for AR0387). The authors reported 76 differentially expressed genes (DEGs) in the in vitro comparison and 259 DEGs in the in vivo comparison, using thresholds of FDR < 0.01 and |LFC| >= 1.0. Our reanalysis achieved strong correlation with the published results. For the in vitro condition, we matched 76 genes with R2 = 0.98 and 100% direction agreement. The in vivo analysis matched all 259 DEGs with R2 = 0.9998 and 100% direction agreement. The key adhesin genes highlighted in the paper showed excellent concordance. SCF1 exhibited LFC of 8.61 (paper) versus 8.67 (our analysis) in vitro, and 4.47 versus 4.53 in vivo. ALS4112 showed similarly close agreement: 5.07 versus 5.08 in vitro, and 2.56 versus 2.56 in vivo.

### Maintaining provenance

Integrating AI agents with analytical platforms like Galaxy presents a provenance challenge. When analysis alternates between Galaxy and external AI-generated scripts, the chain of reproducibility breaks—Galaxy cannot track code executed outside its environment, and AI agents generate numerous artifacts (Python scripts, intermediate files) that are difficult to document systematically. To preserve provenance, we configured our AI agent to invoke Galaxy’s native tools through the API rather than manipulating data directly (this was done via CCA “command” concept; see the following rule). When a CCA interacts with Galaxy via API, it can directly manipulate datasets and collections—but this bypasses Galaxy’s tool framework, losing reproducibility and workflow compatibility. By constraining the agent to use Galaxy tools (e.g., Apply Rules, Filter, `DESeq2`), all operations remain tracked in Galaxy histories, can be extracted into reusable workflows, and produce identical results on re-execution. However, some operations—such as comparing our results against published data—require custom code that Galaxy does not natively support. To address this gap, we developed a standalone JupyterLite notebook (with the help of the same CCA) that reproduces the validation figures shown here without any AI involvement (notebooks can be accessed in Galaxy histories [22,23]). The notebook requires two inputs: (1) `DESeq2` output from Galaxy (TSV with Gene_ID, log2FoldChange, padj columns) and (2) publication data reformatted as CSV with gene_id and log2fc columns. In our workflow, the AI agent’s role was limited to extracting and reformatting data from Excel supplementary files into this simple CSV structure—a step that becomes unnecessary when published supplementary data already conforms to standard formats. Ideally, the CCA functionality should be tightly integrated with BRC-Analytics and Galaxy—a current development priority for us (see below).

## Discussion

### Toward standardization of fungal genomics

Both studies achieved strong validation status, with Santana et al. showing R^2^ = 0.89-0.94 across comparisons and Wang et al. achieving R^2^ = 0.98-0.9998 in both conditions. These results demonstrate that Galaxy-based reanalysis using standard workflows produces results highly consistent with published analyses, and that differential expression patterns are reproducible when using the same statistical methods and significance thresholds.

### AI mistakes and the importance of validation

Our analysis provides a cautionary tale about AI-assisted research. Initially, Claude Code Agent proposed an alternative approach to gene ID mapping: matching genes between annotation versions by finding those with the most similar log2 fold-change values. To the untrained eye this suggestion sounded “scientific”. This LFC correlation method appeared remarkably successful—when we plotted published versus our fold changes, we obtained R^2^ = 0.9996, suggesting near-perfect correspondence. However, subsequent comparison against the official NCBI `old_locus_tag` mapping revealed that only 1% (2 of 203) of LFC-matched gene pairs were correct. The high R2 was an artifact: genes with coincidentally similar fold changes were matched, not the same genes. This example illustrates a fundamental limitation of AI systems: they can propose plausible methods that produce convincing-looking results while being fundamentally flawed. The error was undetectable from the output alone—only independent validation against authoritative sources (NCBI official mappings, confirmed by protein sequence identity) revealed the problem. We recommend that researchers using AI assistants always validate outputs against independent sources, treat high statistical agreement with appropriate skepticism when methodology is novel, and prioritize authoritative reference data over heuristic approximations.

### Importance of LLMs and their responsible use

For reproducibility, LLM-assisted analyses should be conducted through agentic coding tools such as Claude Code or Gemini Code Assist rather than chat-based interfaces. These tools automatically track all generated artifacts—scripts, intermediate files, and analysis outputs—within version-controlled repositories (e.g., GitHub), creating a complete audit trail. While this workflow may currently seem complex for bench biologists unfamiliar with command-line interfaces and version control, it represents the future of data analytics in biology. The interfaces will evolve: emerging tools like Claude Code Web promise to deliver agentic capabilities through browser-based environments, lowering the barrier to entry while maintaining full provenance tracking. As these tools mature, the combination of natural language interaction and automatic versioning will make reproducible AI-assisted analysis accessible to researchers regardless of their computational background.

All results produced with agentic AI tools require independent validation. In this study, we had the advantage of known expected outcomes—published results against which to benchmark our AI-assisted reanalysis. This “ground truth” allowed us to confirm that the AI-directed workflow produced biologically accurate results. However, for novel research where expected outcomes are unknown, researchers must exercise heightened scrutiny. AI agents can confidently produce plausible but incorrect interpretations, and without validation benchmarks, such errors may go undetected. We recommend orthogonal validation approaches: qRT-PCR confirmation of key findings, biological replication, functional studies, and cross-referencing with independent datasets. The provenance tracking enabled by agentic tools becomes especially valuable here—complete audit trails allow retrospective verification when questions arise about specific analytical decisions.

### Why BRC-Analytics/Galaxy for AI-assisted analysis

The choice of workflow platform significantly impacts the feasibility of AI agent integration. For example, Galaxy’s architecture offers distinct advantages over code-first systems like Nextflow for agentic AI workflows. Galaxy provides structured tool metadata through repositories like IUC (github.com/galaxyproject/tools-iuc), where each tool’s parameters, input/output types, and documentation are defined in machine-readable XML. This allows AI agents to query available tools, understand valid parameter options, and make informed decisions—capabilities that require parsing documentation or source code in Nextflow. Galaxy’s stateful API enables agents to inspect histories, monitor job status, and retrieve results through structured endpoints, whereas Nextflow requires log parsing and manual file path management. Perhaps most importantly, Galaxy’s integration with ACCESS-CI provides free, zero-configuration access to high-performance computing, eliminating the infrastructure barriers (container configuration, HPC authentication, resource allocation) that Nextflow imposes on users. Additionally, ACCESS-CI provides access to open LLMs, which would enable functionality similar to the shown here but at no cost to the user.

These architectural differences have practical implications for democratizing AI-assisted genomics. Galaxy’s web-based interface means users need only a browser, while AI agents handle the complexity of tool selection and parameter configuration through the API. Nextflow’s code-generation paradigm, while flexible, requires users to review generated DSL2 scripts, configure execution environments, and debug failures—skills that remain barriers for bench biologists. As AI agents become integral to computational biology, platforms that provide structured metadata, stateful APIs, and accessible infrastructure will enable broader adoption than those requiring programming expertise to operate.

However, capable AI agents partially flatten these distinctions. Tool discovery, parameter selection, log parsing, and error diagnosis—tasks that once separated user-friendly platforms from code-centric ones—become the agent’s responsibility regardless of backend. Once researchers adopt CLI-based AI tools, the barrier to Nextflow drops as well. The more substantial advantages emerge when AI is integrated directly into the platform (see below).

### Agentic AI on-board

We envision tighter integration between BRC-Analytics, Galaxy, and agentic AI systems. Currently, our workflow requires manual coordination: launching analyses through BRC-Analytics, managing data in Galaxy histories, and directing AI agents via API calls. Future development will embed AI agents directly within the Galaxy interface, enabling researchers to describe analyses in natural language while the system automatically selects appropriate workflows, configures parameters, and interprets results. This approach carries higher implementation risk but offers distinct advantages unavailable to external agents. Users require no model setup, API credentials, or payment—the AI is simply part of the browser interface. Agent actions become first-class objects tied to histories, datasets, and provenance records, making AI decisions auditable and inspectable. Combined with Galaxy’s existing reproducibility infrastructure and ACCESS-CI compute, this achieves near-maximal practical reproducibility for LLM-assisted analysis: every prompt, tool invocation, and result lives within a single traceable environment. For researchers without programming backgrounds, integrated AI removes the external tooling barrier entirely—no terminal, no configuration, no credential management. This integration path represents our development priority (Figure 1B).

## Materials and Methods

## Literature Survey and Data Source Identification

To quantify *C. auris* sequencing data, we analyzed complete NCBI SRA database for taxonomy ID 498019 (*Candidozyma auris*) accessed December 3, 2025. SRA metadata (Cauris_SRA.csv) contained 27,201 total runs across 237 BioProjects. RNA-seq represents 812 runs (3.0%) and 64 BioProjects (27.0%), with WGS dominating run counts (26,201 runs, 96.3%) but representing 168 BioProjects (70.9%). Average runs per project: RNA-seq 12.7, WGS 156.0.

To characterize methodology across published RNA-seq studies, we linked all 64 RNA-seq Bio-Projects to associated publications. For each BioProject, we queried EuropePMC REST API (https://www.ebi.ac.uk/europepmc/webservices/rest/) for papers mentioning BioProject accession in full text, and NCBI E-utilities (elink.fcgi) for direct BioProject-to-PubMed links. This identified 21 papers linked to 20 of 64 BioProjects (31%); 44 BioProjects had no linked publications (unpublished or preprint). For papers with PMC IDs (17/20), we retrieved full-text XML and extracted reference genome information by pattern matching (GenBank/RefSeq accessions, strain names, clade designations) and RNA-seq tools (aligners, quantification tools, DE packages). Results in Supplementary Table 2.

For re-analysis validation, we selected Santana et al. (2023) *Science* (PRJNA904261) [20] and Wang et al. (2024) *Nature Communications* (PRJNA1086003) [21].

### WGS Data Contributor Analysis

To characterize sources of *C. auris* WGS data, we analyzed the “Center Name” field from SRA metadata for all 26,201 WGS runs. Organization names were extracted and aggregated by run count and unique BioProjects. Abbreviated center names were expanded using geographic location metadata (geo_loc_name field) to disambiguate state-level public health laboratories (e.g., “MDH_CSL” mapped to Maryland via “USA:Mid-Atlantic” region; “NSPHL” mapped to Nevada via “USA:Nevada” location). Organizations were categorized into: US State/Local Public Health Laboratories, CDC, International Public Health agencies, Academic/Research institutions, and Other. Results presented in Supplementary Table 1.

### Counting Workflow

All analyses used *Candidozyma auris* B8441 reference genome GCA_002759435.3 obtained BRC-Analytics (which mirrors NCBI Datasets). We then use an RNA-seq analysis workflow for obtaining gene counts [24] (Figure 5). For paired-end data, the workflow begins with `fastp` for adapter removal and quality filtering, discarding reads shorter than 15 bp. Filtered reads are aligned to the reference genome using `STAR` with ENCODE-standard parameters, which simultaneously generates gene-level counts. Quality metrics from all steps are aggregated by `MultiQC` into a comprehensive report. The workflow also generates strand-specific coverage tracks (bigWig format) for genome browser visualization. All tools, versions, and parameters are locked within the workflow definition, ensuring identical results across executions.

**Figure 5.**
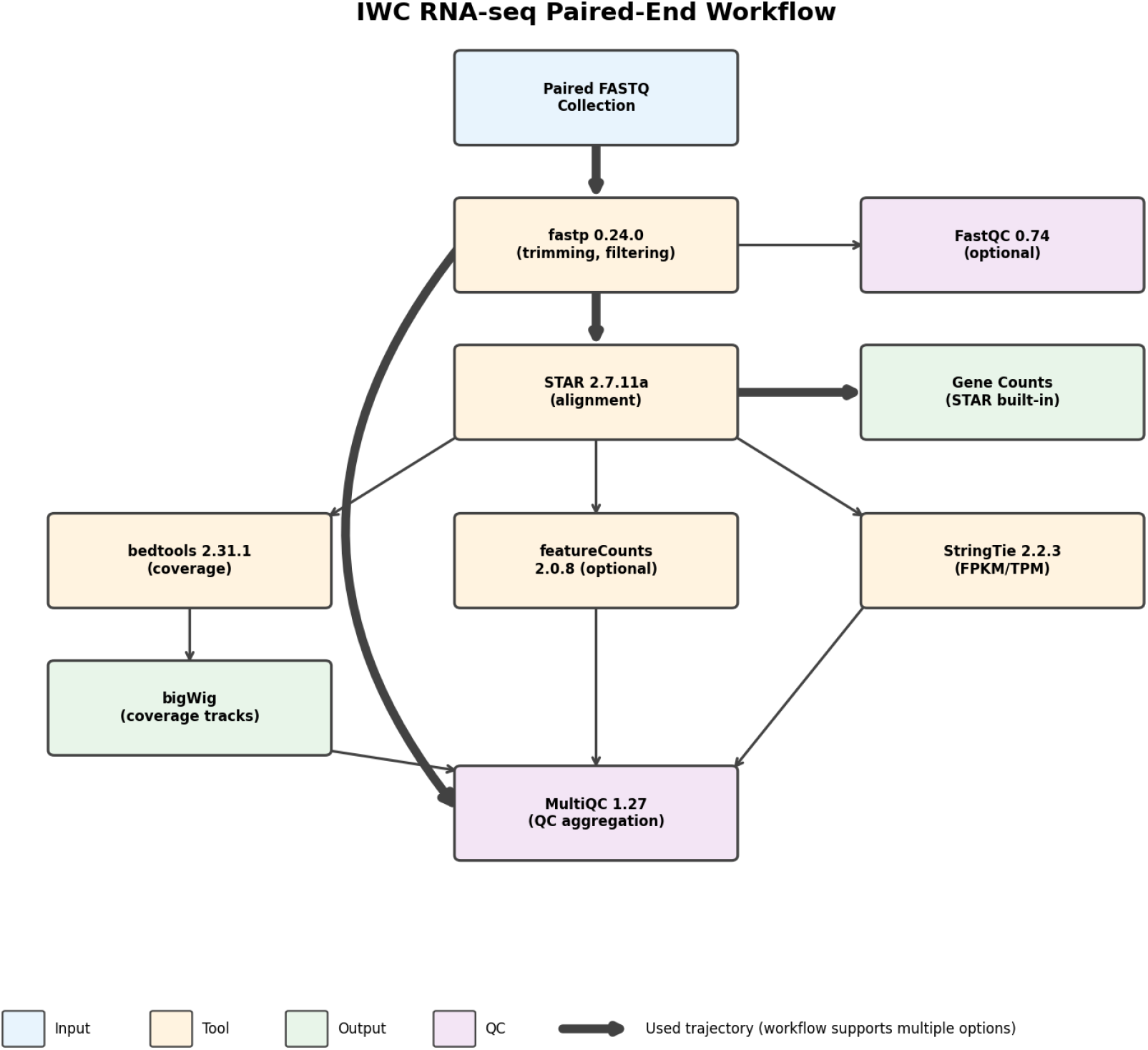
IWC paired-end RNA-seq workflow. The pipeline processes FASTQ files through quality filtering (`fastp`), alignment (`STAR`), and quantification, with optional coverage track generation and QC aggregation via `MultiQC`.

### Differential Expression Analysis

Gene count matrices from `STAR` were analyzed using `DESeq2` (v2.11.40.8+galaxy0) through Galaxy interface. For **Santana et al. dataset**: Samples organized into three collections (AR0382 n=2, AR0387 n=2, tnSWI1 n=2). Two pairwise comparisons performed: (1) AR0382 vs tnSWI1, (2) AR0382 vs AR0387. For **Wang et al. dataset**: Samples split into four collections by strain and condition (AR0382 *in vitro* n=3, AR0387 *in vitro* n=3, AR0382 *in vivo* n=3, AR0387 *in vivo* n=4). Two pairwise comparisons performed: AR0382 vs AR0387 in (1) *in vitro* and (2) *in vivo* conditions.

`DESeq2`parameters: size factor normalization, Benjamini-Hochberg FDR correction, significance threshold FDR less than 0.01, fold change absolute value of log2FC greater than or equal to 1 for Wang dataset. Default parameters used for Santana dataset to match published analysis.

### Gene Annotation Mapping

Published papers used older B8441 annotation versions (GCA_002759435.2) with 6-digit gene ID suffixes (e.g., B9J08_001458) while our analysis used GCA_002759435.3 with 5-digit suffixes (e.g., B9J08_03708). To reconcile gene identities, we used the official NCBI `old_locus_tag` attribute present in the GCA_002759435.3 GTF annotation file, which provides authoritative correspondence between annotation versions. We validated this mapping by extracting protein sequences for mapped gene pairs from both annotation versions and confirming 100% sequence identity. Mapping quality assessed using Pearson correlation, R2, direction agreement percentage, and mean LFC difference. An initial AI-proposed approach using LFC correlation (matching genes by similar fold-change values) was abandoned after validation showed only 1% accuracy despite apparent R2 = 0.9996 (see Discussion). This analysis is saved as a JupyterLite notebook in Galaxy history associated with each paper (see below).

### AI agent integration setup

Claude Code Agent (CCA) interacts with Galaxy through its REST API using an API key stored as an environment variable (`GALAXY_API_KEY` ). The key grants CCA permission to create histories, upload data, execute tools, and retrieve results on behalf of the user. API keys are generated through Galaxy’s user preferences and never committed to version control.

To ensure reproducibility, we configured CCA to prefer Galaxy’s native tools over direct API manipulation. While CCA can programmatically create or modify Galaxy collections via API calls, such operations bypass Galaxy’s tool framework—losing provenance tracking and preventing workflow extraction. Instead, project-level instructions direct CCA to use Galaxy’s built-in collection tools (e.g., `FILTER_FROM_FILE, RELABEL_FROM_FILE, APPLY_RULES` ) for all data transformations. A custom slash command (`/galaxy-transform-collection;` https://github.com/jmchilton/galaxy-agentic-collection-transform) provides CCA with detailed documentation of 26+ collection manipulation tools and decision frameworks for selecting appropriate operations. This approach ensures every analytical step appears in Galaxy’s history as a tool invocation with full parameter capture, enabling complete workflow reconstruction.

### Galaxy Workflows and Reproducibility

All analyses performed on Galaxy Main server (https://usegalaxy.org). Galaxy histories containing complete analysis workflows, intermediate files, and final results are publicly accessible: - Santana et al.: https://usegalaxy.org/u/cartman/h/prjna904261-perm - Wang et al. (Analysis): https://usegalaxy.org/u/cartman/h/prjna1086003-perm

These histories also contain the JupyterLite notebooks used for validation analysis and figure generation.

IWC workflows used are available at https://iwc.galaxyproject.org and are version-controlled in GitHub repository at https://github.com/galaxyproject/iwc. Workflow diagrams and analysis reports available in supplementary materials.

## Acknowledgements

We would like to express our immense gratitude to Dan Stanzione and David Hancock for essential computational resources provided by the Advanced Cyberinfrastructure Coordination Ecosystem (ACCESS-CI), Texas Advanced Computing Center, and the JetStream2 scientific cloud. This work is funded by the NIH Grant U24AI183870.

## Supplementary Materials

**Supplementary Table 1:** *C. auris* WGS data contributors by organization category and top sequencing centers.

*Panel A: Summary by Organization Category*
| Category | Organizations | Runs | % of Total |
| --- | --- | --- | --- |
| US State/Local Public Health Labs | 26 | 20,552 | 78.4% |
| CDC | 2 | 2,626 | 10.0% |
| Academic/Research | 46 | 1,365 | 5.2% |
| Other | 41 | 1,345 | 5.1% |
| International Public Health | 5 | 313 | 1.2% |
| **TOTAL** | **120** | **26,201** | **100%** |

*Panel B: Top 15 Contributing Organizations*
| Organization | Full Name | Runs | % |
| --- | --- | --- | --- |
| UPHL_ID | Utah Public Health Laboratory | 4,447 | 17.0% |
| NVSPHL | Nevada State Public Health Laboratory | 4,363 | 16.7% |
| CDC-NCEZID-MDB | CDC Mycotic Diseases Branch | 2,406 | 9.2% |
| MDH_CSL | Maryland Dept of Health, Central Services Lab | 2,309 | 8.8% |
| TXDSHS | Texas Dept of State Health Services | 1,487 | 5.7% |
| MDHHS-GS | Michigan Dept of Health & Human Services | 1,289 | 4.9% |
| - | Wisconsin State Laboratory of Hygiene | 1,211 | 4.6% |
| RIPHL | Rhode Island Public Health Laboratory | 1,197 | 4.6% |
| NSPHL | Nevada State Public Health Laboratory | 1,031 | 3.9% |
| - | Wadsworth Center (New York) | 705 | 2.7% |
| - | Minnesota Dept of Health | 688 | 2.6% |
| OCPHL_CA | Orange County Public Health Lab (California) | 659 | 2.5% |
| - | Washington State Dept of Health | 583 | 2.2% |
| UNLV NPM | Univ of Nevada Las Vegas, Pathogen Monitoring | 443 | 1.7% |
| - | Fudan University | 264 | 1.0% |

*US public health laboratories (state/local + CDC) account for 88.4% of all C. auris WGS data, reflecting outbreak surveillance priorities. Nevada appears twice (NVSPHL + NSPHL = 5,394 runs, 20.6%), indicating major outbreak focus.*

**Supplementary Table 2:** RNA-seq methodology across 20 published *C. auris* BioProjects with linked publications (2018-2025).

| BioProject | PMID | Authors | Year | Runs | Reference Genome | RNA-seq Tools |
| --- | --- | --- | --- | --- | --- | --- |
| PRJNA445471 | 30559369 | Muñoz JF et al. | 2018 | 24 | B8441, B11220, B11243 | Bowtie2, TopHat2, RSEM, Trinity, edgeR |
| PRJNA477447 | 29997121 | Kean R et al. | 2018 | 22 | B8441 (de novo) | Trinity, HISAT2, Kallisto, DESeq2 |
| PRJNA682185 | 34630944 | Zamith-Miranda D et al. | 2021 | 36 | B8441 (GCA_002759435.2) | DESeq2, edgeR |
| PRJNA682422 | 34180774 | Lara-Aguilar V et al. | 2021 | 6 | B8441 (GCA_002759435.2) | FastQC, Trimmomatic, fastp, STAR, featureCounts, DESeq2 |
| PRJNA735406 | 34354695 | Zhou W et al. | 2021 | 6 | B11221 (Clades I-V) | Trimmomatic, HISAT2, Cufflinks, HTSeq, DESeq2 |
| PRJNA801628 | 35473297 | Biermann AR et al. | 2022 | 24 | B8441, B11221, B11243 (Clades I, III, IV) | HISAT2, featureCounts, edgeR |
| PRJNA830685 | 36445083 | Narayanan A et al. | 2022 | 16 | B8441, CBS10913 (Clade II) | FastQC, fastp, BWA, Bowtie2, HTSeq, DESeq2 |
| PRJNA788930 | 35652307 | Shivarathri R et al. | 2022 | 12 | NS | RNA-seq |
| PRJNA792028 | 36913408 | Bing J et al. | 2023 | 15 | GCA_002759435.2, GCF_002775015.1 | HiSat2, StringTie, DESeq2, BWA |
| PRJNA904261 | 37769084 | Santana DJ et al. | 2023 | 6 | B8441 (Clade I) | RNA-seq |
| PRJNA1015296 | 38493178 | Bing J et al. | 2024 | 141 | B8441 (GCA_002759435.2) | HiSat2, StringTie, DESeq2, BWA |
| PRJNA902676 | 38722168 | Yang B et al. | 2024 | 40 | B11220, B11221 (Clades II, III) | Kallisto, DESeq2 |
| PRJNA1036037 | 39480072 | Li J et al. | 2024 | 22 | Clade IV | RNA-seq |
| PRJNA1086003 | 39455573 | Wang TW et al. | 2024 | 13 | B8441 (Clade I) | HISAT2, STAR, DESeq2 |
| PRJEB57846 | 39297640 | Rhodes J et al. | 2024 | 12 | NS | WGS, RNA-seq |
| PRJNA1012821 | 40468551 | Chauhan A et al. | 2025 | 16 | B8441, B11220 (CGD) | FastQC, fastp, Bowtie2, HTSeq, DESeq2 |
| PRJNA1139166 | 40099908 | Phan-Canh T et al. | 2025 | 15 | B8441 (GCA_002759435.2) | FastQC, fastp, cutadapt, STAR, featureCounts |
| PRJNA1208975 | 40530673 | Yang G et al. | 2025 | 9 | Clade I | RNA-seq |
| PRJNA1232830 | 40066990 | Chauhan M et al. | 2025 | 6 | Clade I | RNA-seq |
| PRJNA1291775 | 40863525 | Vidal-Montiel A et al. | 2025 | 6 | GCA_003014415.1, GCA_034640365.1 (Clades III, IV) | FastQC, Trimmomatic, STAR, featureCounts, DESeq2 |

*NS = Not specified in available text. Data extracted from PMC full text via EuropePMC and NCBI E-utilities APIs. 44 additional BioProjects had no linked publications.*

### Supplement 1

A “plan” generated by Claude Code:

Goal: Split collection #244 into 3 condition-specific collections for `DESeq2`. Experimental conditions (from paper):

| Condition | Description | Replicates |
| --- | --- | --- |
| AR0382_WT | Clade I reference, highly adhesive | 2 (A, B) |
| AR0387_WT | Clade I, poorly adhesive (206 SNPs from AR0382) | 2 (A, B) |
| tnSWI1 | SWI1 insertional mutant of AR0382 | 2 (A, B) |

Expected SRR mapping (to verify): - AR0382_WT: SRR22376031, SRR22376032 - AR0387_WT: SRR22376029, SRR22376030 - tnSWI1: SRR22376027, SRR22376028

Approach (from galaxy-agentic-collection-transform): 1. Fetch SRA metadata → confirm mapping 2. Check collection #244 element naming 3. Create identifier files for each condition 4. Use **FILTER_FROM_FILE** Galaxy tool to split 5. Rename output collections

Next steps before execution: 1. Verify SRA metadata 2. Check collection #244 element names 3. Present findings for your approval

